# Automated video-based heart rate tracking for the anesthetized and behaving monkey

**DOI:** 10.1101/2020.06.23.167411

**Authors:** Mathilda Froesel, Quentin Goudard, Marc Hauser, Maëva Gacoin, Suliann Ben Hamed

## Abstract

**Background:** Heart rate is extremely valuable in the study of complex behaviours and their physiological correlates in non-human primates. However, collecting this information is often challenging, involving either invasive implants or tedious behavioural training.

**New Method:** In the present study, we implement a Eulerian Video Magnification (EVM) heart tracking method in the macaque monkey combined with wavelet transform. This is based on a measure of image to image fluctuations in skin reflectance due to changes in blood influx.

**Results:** We show a strong temporal coherence and amplitude match between EVM-based heart tracking and ground truth ECG, from both color (RGB) and infrared (IR) videos, in anesthetized macaques, to a level comparable to what can be achieved in humans. We further show that this method allows to identify consistent heart rate changes following the presentation of conspecific emotional voices or faces.

**Comparison with Existing Method(s):** Eulerian Video Magnification (EVM) is used to extract heart rate in humans but has never been applied to non-human primates. Video photoplethysmography allows to extract awake macaques heart rate from RGB videos. In contrast, our method allows to extract awake macaques heart rate from both RGB and IR videos and is particularly resilient to the head motion that can be observed in awake behaving monkeys.

**Conclusions:** Overall, we believe that this method can be generalized as a tool to track heart rate of the awake behaving monkey, for ethological, behavioural, neuroscience or welfare purposes.

**Highlights:**

- Heart rate varies during complex non-human primate (NHP) behaviour and cognition.
- We apply Eulerian Video Magnification to track NHP heart rate (EVM-HR).
- EVM-HR can be used with RGB & IR videos, and anesthetized or awake NHPs.
- NHP EVM-HR vary with emotional content of presented stimuli.
- EVM-HR is of interest to ethology, behavioural, neuroscience & welfare purposes.

## Introduction

Tracking variations in autonomous responses has proven to be invaluable in the study of complex behaviours and their physiological correlates in non-human primates (Bradley et al., 2017). These include tracking changes in pupil diameter (Bradley et al., 2008; Henderson et al., 2018; Macatee et al., 2017; Reynaud et al., 2019), in skin conductance (Nakasone et al., 2005), social blinks (Ballesta et al., 2016), blink rates (Cléry et al., 2017; Guipponi et al., 2015), nose temperature (Kuraoka & Nakamura, 2011) and heart rate (Grandi & Ishida, 2015; Hassimoto et al., 2002). Heart rate measure is of particular relevance in diverse cognitive contexts. For example, it has been shown that heart rate increases when monkeys watch videos with high affective content (Bliss-Moreau et al., 2013) and during learning process (Uchiyama, Ohtani, and Ohta 2007, Mitz et al. 2017). In spite of this, very few methods currently allow to easily, reliably and non-invasively track heart rate in awake behaving untrained monkeys.

Several tools are already available on the market to extract heart rate, such as electrocardiograms (ECG) or pulse oximeters. ECG detects changes in voltage generated by the cardiac muscles and requires to place electrodes on a shaved skin. Pulse oximetry measures oxygen blood saturation and blood pulsations thanks to photoplethysmography (PPG). PPG consists in detecting luminosity variations of the skin that are directly related to changes in blood flow (Challoner & Ramsay, 1974; Kamal et al., 1989). It involves placing a captor on the subject, usually on the fingers. For both these methods, signal quality can deteriorate in time, due to a displacement of the electrodes or the captor. In addition, when dealing with adult human subjects, this may introduce experimental biases due to the fact that subjects possibly becomes aware of the scope of the study. In young infants or animals, placing the captor or maintaining it all throughout the experiment may turn out to be challenging if not impossible. In monkeys, recording the heart rate either involves ECG or PPG under sedation (Huss et al., 2015; X. Sun et al., 2013; Yamaoka et al., 2013) or implanting a telemetry device for heart rate measure during behaviour (Chui et al., 2012; Hoffmann et al., 2014). PPG recording or heart rate measures using captors embedded in a wearable jacket is also an option (Derakhchan et al., 2014; Hassimoto et al., 2002; Kremer et al., 2011). However, this requires intensive monkey training and might bias heart rate measures due to discomfort or stress.

Recently, methods allowing to extract heart rate at a distance from human subjects have been developed based on video image processing (Alghoul et al., 2017; Madan et al., 2018; WuHao-Yu et al., 2012). Imaging photoplethysmogram (iPPG), detects, as is the case for PPG, variation of skin light absorption/reflection properties (Huelsbusch & Blazek, 2002; Takano & Ohta, 2007; Verkruysse et al., 2008). Indeed, heartbeat induces blood flow in all the body including the face skin. This results in a change in the skin reflectance (Anderson & Parrish, 1981; Angelopoulou, 2001; Jakovels et al., 2013, 2012; Tsumura et al., 2003). These changes in human skin reflectance can be tracked from webcam (Poh et al., 2010, 2011) or smartphone (Kwon et al., 2012) video quality images associated with ICA signal processing techniques.

These methods appear highly relevant to non-human primate research and welfare, as cognitive processes, emotional states or welfare indicators such as stress can only be inferred by indirect measures. However, one of the major challenges in this context is the fact that changes in skin reflectance are more difficult to detect in monkeys due reduced glabrous facial skin surface. In spite of this limitation, Unakafov et al. (2018) have successfully applied heart rate tracking in awake macaques in combination with discrete Fourier and wavelet transform based iPPG.

Here, we present an alternative heart rate tracking method in the monkey, using Eulerian Video Magnification (EVM) (WuHao-Yu et al., 2012). A major advantage of this approach relative to iPPG is its resilience to subject motion (Alghoul et al., 2017). We have further associated EVM video extraction with wavelet transformation based analyses, that has also been shown to be motion tolerant on poor quality human webcam video data (Bousefsaf et al., 2013). In a first step, we show that EVM-based heart rate tracking has a high temporal coherence with ground truth ECG data, whether extracted from RGB video images or IR video images (that are often used in neuroscience experimental protocols) and that EVM-based heart rate estimate is very close to ECG-based heart rate estimate. For both types of video quality, we show that temporal coherence between EVM-based heart tracking and ground truth ECG is not significantly different between humans and monkeys. Last, we describe the dependence of temporal coherence between EVM-based heart tracking and ground truth ECG on the localization of the specific facial region EVM is performed onto. In a second step, we apply EVM-based heart tracking to the awake monkey and we show that this measure allows to identify consistent heart rate changes following the presentation of conspecific emotional voices or faces. Overall, we believe that this method can be generalized as a tool to track heart rate of the awake behaving monkey, for ethological, behavioural or welfare purposes.

## Material and methods

### Subjects

#### Monkeys

Three rhesus monkeys (*Macaca mulatta*) participated at this study. They were aged between 9 and 17 years (2 males: monkeys T and S, 1 female: monkey Z). The project was authorized by the French Ministry for Higher Education and Research (project no. 2016120910476056 and 2015090114042892) in agreement with the French implementation of European Directive 2010/63/UE. This authorization was based on the ethical evaluation by the local Committee on the Ethics of Experiments in Animals (C2EA) CELYNE registered at the national level as C2EA number 42 and the National Ethics of Experiments in Animals board of the French Minsitry of Higher Education, Research and Innovation.

#### Humans

Two human participants were included in this study and were covered by a broader project authorization (ID RCB 2018-A03438-47).

### Monkey anaesthesia

Monkeys were lightly anesthetized with Zoletil (Tiletamine-Zolazepam, Virbac, 10 mg/kg) so as to avoid head movements. During the video and ECG acquisitions, monkeys were gently resting on their side, under constant physiological monitoring.

### Monkey Behavioural task

During these sessions, monkeys were awake and sat in a sphinx position in a plastic monkey chair and head restrained to avoid movement. They faced a screen (1920×1200 pixels and a refresh rate of 60 Hz) placed 54 cm from their eyes on which stimuli were presented by an experimental control and stimulus delivery software (EventIDE^®^). The auditory stimuli were displayed by a Sensimetrics^®^ S14 insert earphones and set up. Monkeys were required to fixate the centre of the screen all throughout the recording blocks. Eye fixation was controlled thanks to an Eyelink video eye tracker (EyeLink^®^). Recording blocks consisted in 2 alternations of 16 seconds of fixation and 16 seconds of emotional stimuli presentation (alternations of 450ms stimuli of the same sensory and emotional category, differing in specific identity). In one type of recording blocks, fixations alternated with highly emotional auditory content (screams). In a second type of recording blocks, fixations alternated with highly emotional visual content (4°x4° aggressive faces). In total, 54 fixation to scream transitions (30 for monkey T and 24 for monkey S), and 57 fixation to aggressive faces (30 for monkey T and 27 for monkey S) were collected. The auditory stimuli and part of the visual stimuli were recorded in Cayo Santiago, Puerto Rico and kindly produced by Marc Hauser. The other visual stimuli were created in our own lab.

### Electrocardiogramm recordings

Electrocadiographic signal (ECG) was recorded thanks to a BiopacSystem^®^. In humans, two electrodes were placed on the subject’s thoracic cage, on each side of the heart, while the reference electrode was placed on the abdomen, close to the stomach.

In the macaque, two electrodes were placed on the subject’s thoracic cage, on each side of the heart, while the reference electrode was placed close to the groin on a shaved skin. ECG signal was recorded at a frequency of 2kHz, so as to have a well-defined QRS waveform.

### Video recordings

We used a USB camera with a variable framerate (maximum frame rate of 30 frames per second), and a spatial resolution of 640×480 pixels. The camera had two working modes depending of light intensity. At high light intensity level, the camera was a color (RGB) device. At low light intensity level, the camera was an infrared (IR) device.

When comparing EVM-based heart rate tracking to ground truth ECG heart rate estimation in monkeys (monkey T and monkey Z), the animals were anesthetized and laying on their side. The video recording was targeted to the face. When comparing EVM-based heart rate tracking to ground truth ECG heart rate estimation in humans (subject H1 and H2), subjects were requested to gently and steadily gaze at the camera and video recording were targeted to the face. For each monkey and human subject, we collect ten minutes of videos in full light (RGB mode) or in the dark (IR mode).

When analysing the effect of emotional stimuli on EVM-based heart rate tracking, we recorded IR videos while monkeys T and S were performing the above described task.

ECG and video time series were synchronized using the AcqKnowledge^®^ software. Specifically, the AcqKnowledge^®^ software was used to record the ECG signal and send start and stop synchronization triggers to the video recording system.

**Figure 1.**
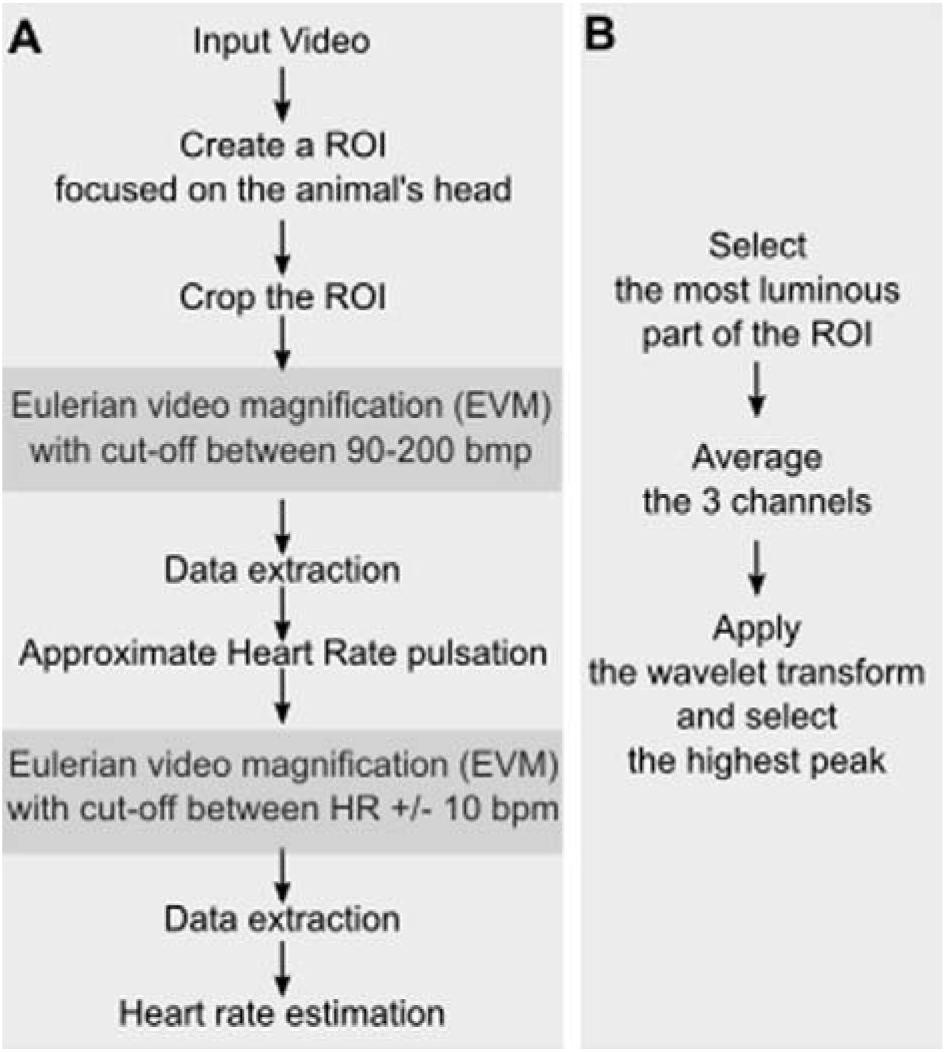
A) ***Heart rate video extraction pipeline***. A Region of interest (ROI) is defined on the video input, ideally placed in a hairless skin region, such as the face. The video is cropped around this ROI selection and serves as input to the Eulerian video magnification (EVM) algorithm (see text for specifications). The result is a video in which luminosity changes due to heart rate are enhanced. A first heart rate (HR) approximation is extracted from this video (as described in B). In order to obtain a more precise HR estimate, this approximated heart rate is used as a parameter for a second EVM processing round. B) ***Heart rate approximation***. Average time series are extracted from the ROI pixels of highest luminosity, for each color channel, and their frequency power profile is extracted using a wavelet transform. Peak frequency is taken as the estimate of HR.

### Heart rate extraction from recorded video

#### Definition of Region of Interest (ROI)

In order to optimize heart rate extraction, the first processing step involves defining, in the recorded video a region of interest (ROI) to feed in the rest of the processing pipeline (figure 1A). Because subsequent processing involves estimating variations in skin luminosity, ROIs should be placed on the face. The optimal location on the face is further discussed in the result section. The video is cropped around this ROI and the output is fed into a first EVM processing step.

#### Eulerian Video Magnification–step one

The Eulerian video magnification algorithm (EVM, WuHao-Yu et al. 2012) allows to magnify variations in frame to frame video information and can thus be used amongst other things to amplify skin colour variations due to blood circulation. It involves both a spatial and a temporal processing such that any variation in pixel properties through time and space is amplified. The input video is decomposed in several spatial frequency bands. The same temporal filter is applied to all these bands. The most relevant spatial frequency band to the signal of interest (here, HR detection) is amplified and the result of this amplification is added to the initial video signal. Amplification factor and temporal filtering parameters are hand-optimized by the experimenters to maximize the identification of the signal of interest. Specifically, we use the EVM functions for Matlab^®^ (WuHao-Yu et al. 2012, http://people.csail.mit.edu/mrub/evm/), and run it on our ROI-cropped video, defining a spatial filtering by a Gaussian blur and a down sample, as well as a temporal filtering by a band pass filter matching the expected range of heart rate estimates (e.g. 90 to 200 bpm; Grandi and Ishida 2015; Tatsumi et al. 1990; Clarke, Mason, and Mendoza 1994). The output of this first EVM processing round is a video in which luminosity changes due to heart rate are enhanced (figure 1A).

#### Heart rate approximation

The ROI pixels with highest frequency power are further extracted, all video colour time series are averaged and a wavelet transform is used to select frequency with highest peak in these selected pixels (figure 1B).

#### Heart rate estimation – step two

In order to obtain a more precise estimate of heart rate in time, the EVM is run onto the selected ROI-cropped video a second time, and a band pass temporal filtering better matching the individual subject heart rate range is applied (figure 1A). A second heart rate estimation is run as described in figure 1B and the Heart rate approximation paragraph above.

### Signal processing

#### ECG-based heart rate

Inter-peak intervals were extracted from the raw ECG signal and pulse rate were estimated in time by computing a running average over the inter-peak times series (figure 2b, averaging window size = 20 s).

**Figure 2.**
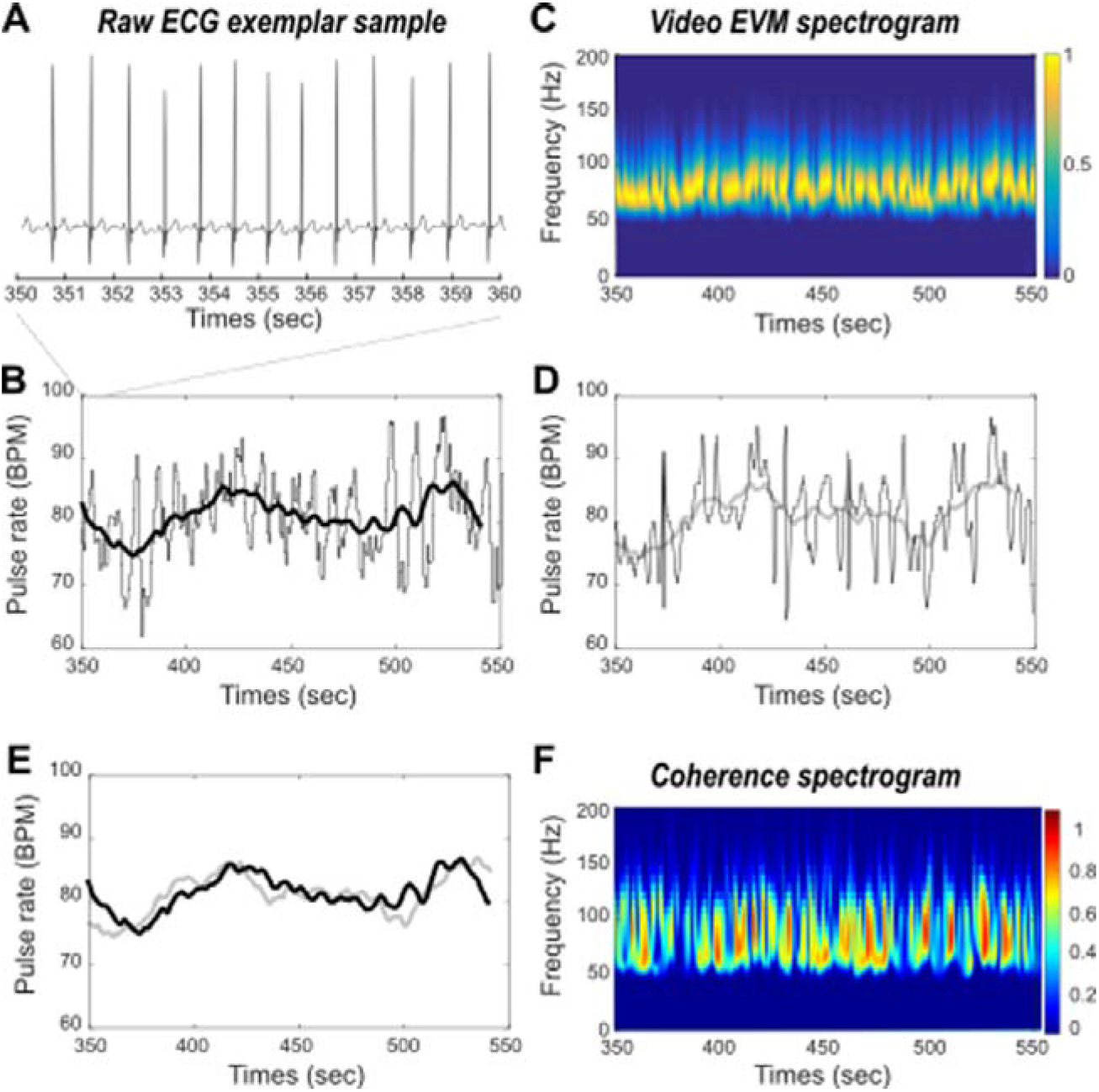
Comparing ECG and EVM heart rate estimates. A) Raw ECG exemplar sample. B) ECG inter-peak interval estimate (gray) and pulse rate running average (black) in time. C) Time frequency spectrogram of EVM processed video data. Highest power frequency band (yellow) corresponds to the heart rate estimate. D) EVM peak frequency estimate (gray) and corresponding pulse rate running average (light gray) in time. E) Overlay of ECG (black) and EVM (light gray) based running average pulse rates estimates in time. F) Coherence between ECG and EVM HR estimates.

#### EVM-based heart rate

Instantaneous heart rate estimates are defined as maximal power frequencies at each time step of the wavelet transform. EVM-based heart rate is estimated in time by computing a running average over the instantaneous heart rate times series (figure 2c, averaging window size = 20 s).

#### ECG – EVM temporal coherence

In order to statistically assess the extent to which ECG-based and EVM-based heart rate measures co-vary in time, we perform a wavelet coherence analysis that estimates instantaneous coherence in time and for all frequencies of interest (figure 3). Reported temporal coherence is the average of maximum temporal coherence in time, across all frequency ranges. The wavelet coherence improves the time-frequency localization of spatial correlation patterns. It is also particularly well suited to non-stationary and noisy physiological data (Chavez & Cazelles, 2019).

**Figure 3.**
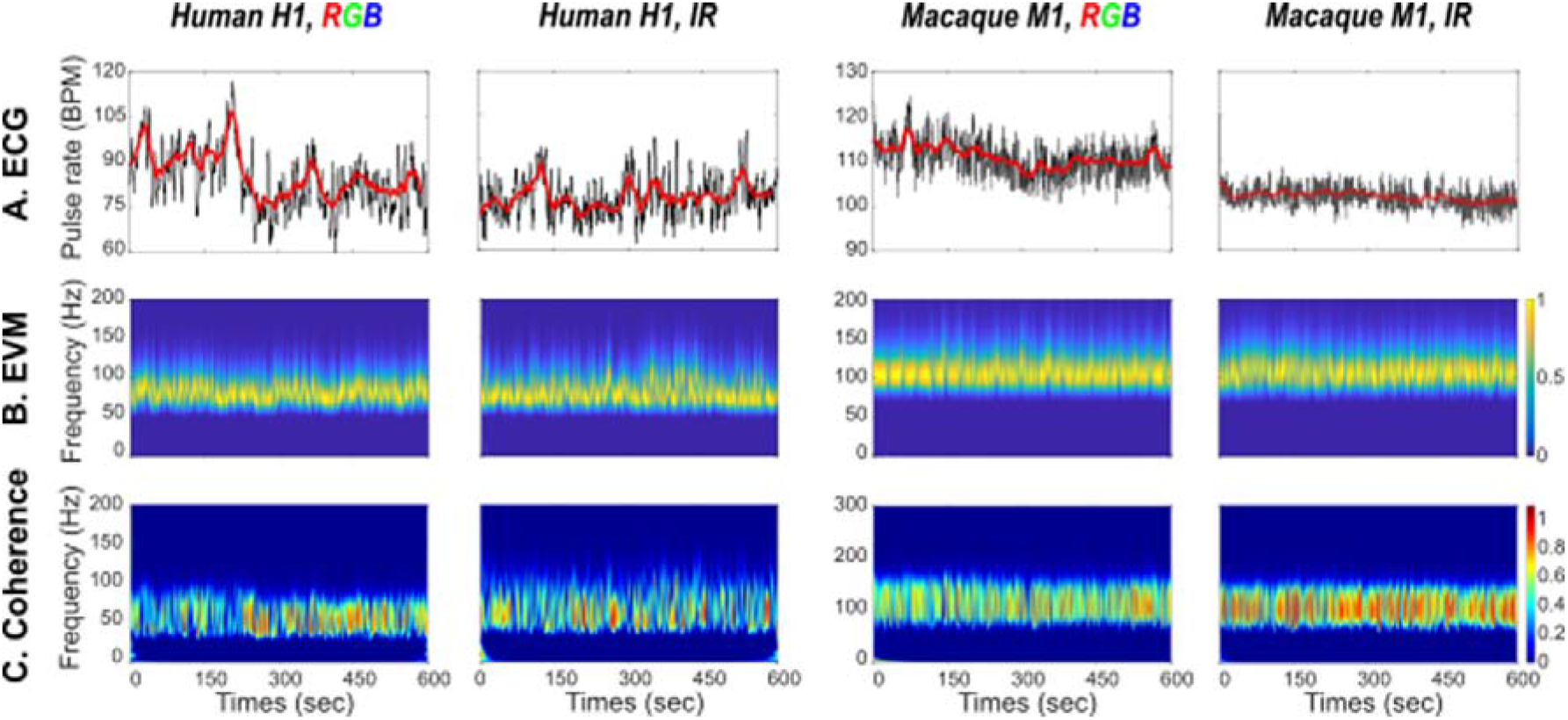
Heart rate estimation in humans (panels 1 & 2) and monkey (panels 3 & 4) based on RGB (panels 1 & 3) and IR (panels 2 & 4). A) ECG HR ground truth. B) EVM HR estimate. C) Coherence between ECG and EVM HR estimates.

#### Stimulus triggered changes in EVM-based heart rate

EVM-based heart rate is estimates for each run and each monkey the pulse rate along the entire recording blocks. Heart rate time series are extracted around the onset of the stimulus of interest (i.e. scream or aggressive face, [-16s 16s]) and realigned to stimulus event presentation (0s time reference). Individual time series are baseline corrected ([−13 −3s]) and mean +/- s.e. is computed in time. Pre ([−13 −3s]) and post-stimulus ([3 13s]) heart rate estimates are compared using a non-parametric Wilcoxon test.

## Results

### Comparing EVM-based heart rate estimation to ground truth ECG

In order to validate the EVM-based heart rate measure, we compared the inter-pulse signal estimated using the EVM approach (figure 1) to ground truth ECG-based inter-pulse estimation. Figure 2A represents a sample of our ground truth recording measure in a human subject, and the corresponding inter-peak estimation (figure 2B, gray) and pulse rate running average on a longer recording period (figure 2B, black). Figure 2C represents, during the exact same time period, the frequency over time, the EVM-based time frequency output of our EVM processing pipeline (figure 1). Heart rate induced rhythmic changes in the recorded video flow are clearly observed in the 70 to 90 Hz frequency range (Figure 2C, yellow saturation epochs). At each time point, EVM-based instantaneous heart rate peak frequencies are extracted resulting in an EVM-based pulse rate estimation in time (Figure 2D, gray, running average, light gray). The ECG-based (Figure 2E, back) and EVM-based running averages (Figure 2E, gray) strongly co-vary in time. In order to statistically assess the extent to which ECG-based and EVM-based heart rate measures co-vary in time, we perform a wavelet coherence analysis that estimates instantaneous coherence in time and for all frequencies of interest (figure 2F). This statistics computes, at each time point of the time series, and each frequency, the degree of coherence between the two signals of interest. On this specific recording, average temporal coherence between ECG-based and EVM-based heart rate estimates is 0.5 (s.d. = 0.22).

### EVM-based heart rate estimation performs better in monkeys than in humans, whether using color (RGB) or infrared (IR) videos

EVM-based heart rate estimation has already been validated with human RGB data (Wu Hao-Yu et al. 2012). Here, we confirm that heart rate can be estimated from human RGB (figure 3, first column) as well as IR videos (figure 3, second column). We further show that EVM-based heart rate estimation can also be achieved from anesthetized macaque RGB (figure 3, third column) as well as IR videos (figure 3, fourth column). For each type of video, each column represents ECG-based inter-pulse estimation and corresponding running average (upper row), normalized EVM-based pulse rate power estimation (middle row) and temporal coherence between ECG-based and EVM-based heart rate estimates (lower row).

Table 1 further summarizes pulse rate statistics for each type of recording. Differences between ECG and EVM-based heart rate estimations range between 0.07 BPM and 8.28 BPM (mean=0.85, s.d.=4.01). Both methods reach very similar signal variability (mean s.d.: ECG: 2.9275; EVM: 2.6575). Average coherence between the two signals is 0.51 (s.d.= 0.08). Importantly, coherence between ECG-based and EVM-based heart rate estimates tend to be higher in monkeys (mean=0.5763, s.d.=0.05) than in humans (mean=0.4554, s.d.=0.05, unilateral Wilcoxon, p=0.06). Overall, this is thus evidence that EVM can be used to provide a reliable non-invasive measure of heart rate estimation in the anesthetized preparation.

**Table 1.** Pulse rate estimation statistics from ECG and EVM time series and corresponding temporal coherence.

|  |  | ECG-based pulse rate |  | EVM-based pulse rate |  | Coherence coefficient |
| --- | --- | --- | --- | --- | --- | --- |
|  |  | mean | s.d. | mean | s.d. |  |
| Human 1 | RGB | 84.10 | 5.64 | 80.83 | 2.9 | 0.48 |
|  | IR | 77.95 | 2.9 | 77.88 | 2.3 | 0.4656 |
| Human 2 | RGB | 84.76 | 4 | 76.48 | 3.3 | 0.377 |
|  | IR | 81.35 | 2.88 | 82.08 | 4.61 | 0.499 |
| Monkey 1 | RGB | 109.5 | 1.74 | 105.79 | 1.49 | 0.554 |
|  | IR | 102.5 | 0.89 | 105.5 | 1.88 | 0.66 |
| Monkey 2 | RGB | 178.35 | 1.94 | 182.26 | 2.95 | 0.5676 |
|  | IR | 167.16 | 3.43 | 168.05 | 1.83 | 0.5236 |

### Effect of ROI selection on EVM-based heart rate estimation

During the EVM signal analysis, a region of interest (ROI) must be defined in order to optimize heart rate extraction. This ROI is usually placed in the most luminous part of the video, in order to record the luminosity variabilities as precisely as possible and thus have the best EVM-based heart rate estimation. (see figure 1B). Although coherence between ECG and EVM-based heart rate estimations is always higher than 0.45, it can reach up to 0.55 when adequately placed on the face. Figure 4A represents the initial video frame recorded in an anesthetized monkey. Figure 4B represents coherence between ECG and EVM-based heart rate estimations computed over 10×10 independent ROIs covering the video frame presented in figure 4A. Coherence is overall higher in the glabrous skin parts of the face (eyes and eye lids, snout and mouth), although important variations can still be observed between different parts of this glabrous skin. The skin around the eyes and the snout appear to be most informative in relation with heart-rate extraction. Focusing on the face regions (figure 4C) and defining 270 smaller pixels in this region produces local pixels of maximal coherence at the bottom of the snout. However, on average, these smaller voxels do not produce higher coherence between ECG and EVM-based heart rate estimations than larger voxels as presented in figure 4B.

**Figure 4.**
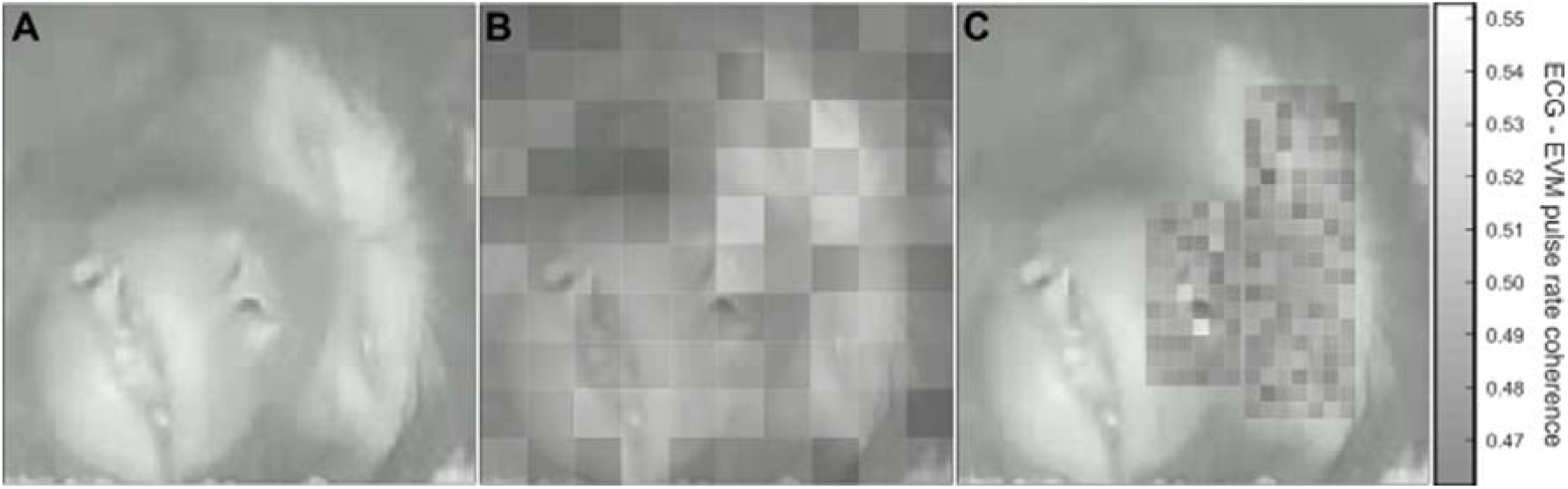
Effect of ROI size and localization on ECG-EVM pulse rate coherence. A) Reference monkey face on which is superimposed ECG-EVM coherence maximum power based on a 10×10 pixel matrix covering the entire face (B) or smaller pixels covering the eyes and muzzle (C). Note that B and C share the same color scale.

### EVM-based heart rate estimation tracks changes in heart rate in the awake behaving monkey

The main objective of this paper is to provide a non-invasive alternative to ECG and pulse oximeter heart rate tracking method for monkeys during typical behavioral tasks, with no behavioral training requirements. During such tasks, heart rate measures have been shown to co-vary with cognitive (Siennicka et al., 2019; Zeki Al Hazzouri Adina et al., 2014) or emotional processes (Bliss-Moreau et al., 2013). A non-invasive heart rate tracking alternative thus needs to be able to track subtle changes in heart rate while monkeys are actively performing a task. Here, we required two monkeys to maintain eye fixation on a central cross for reward, while stimulating them with either monkey screams (figure 5A, auditory negative emotion) or with a static aggressive monkey picture (figure 5B, visual negative emotion). Meanwhile, we recorded their faces thanks to an IR video system. The videos were EVM-processed as described in figure 1, placing a ROI between the two eyes, at the base of the snout, allowing to obtain an optimal estimate of heart rate. Then, we synchronized the latter with events, characterized by the sensory stimuli and corrected it with respect to the pre-event baseline (baseline corrected), such thus we could report the heart rate changes induced by stimulus presentation, irrespective of other possible modulatory effects. Both the monkey screams (figure 5A, pre-post comparison, Wilcoxon p=0.03) and the aggressive monkey faces (figure 5B, p=0.001) induce a small but systematic change in heart rate. The onset of these systematic changes are of the order of a few seconds and are compatible with the reported latency of heart rate changes during emotional processing (Bliss-Moreau et al., 2013).

**Figure 5.**
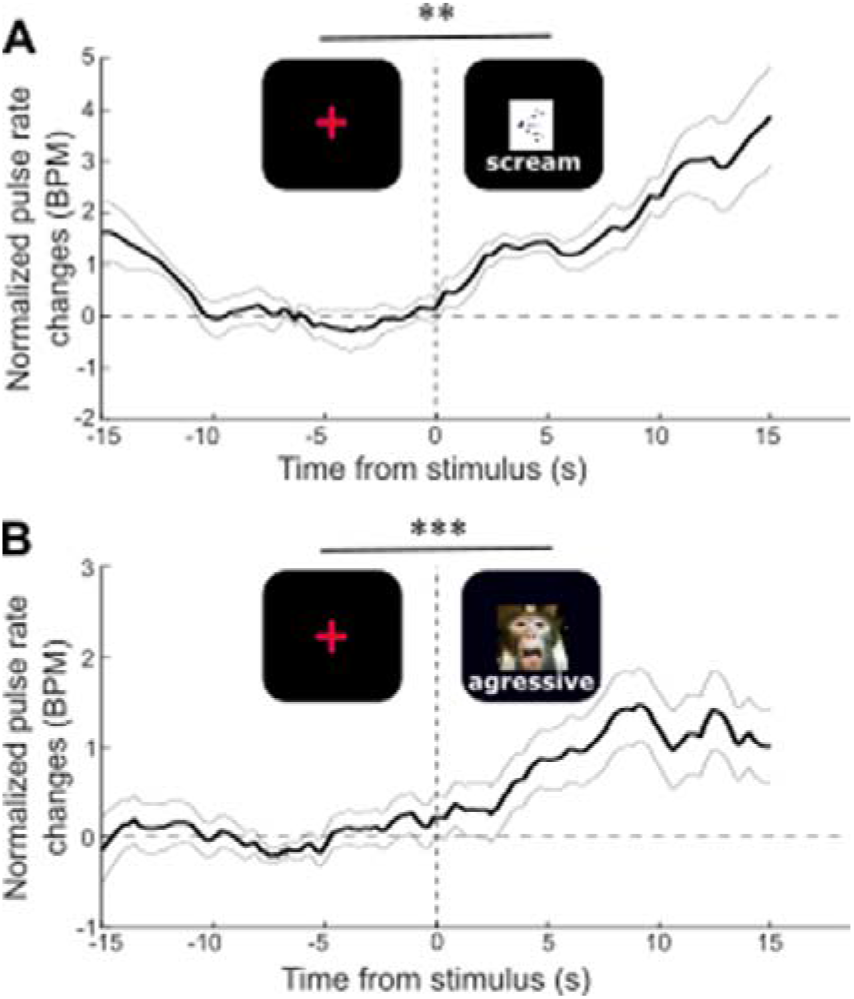
EVM heart rate estimate modulation (mean +/- s.e.) by (A) monkey screams (Wilcoxon test comparing pre-stimulus [−400 −100ms] and post-stimulus [100 400] epochs, p=0.03) and (B) monkey aggressive faces (p=0.001).

## Discussion

Overall, we demonstrate that the Eulerian Video Magnification combined with wavelet transform analyses allows to reliably extract heart rate estimates from both RGB and IR videos of both anesthetized and awake macaque monkeys. These heart rate estimates are directly comparable to ground truth ECG heart rate estimate, as they have the same temporal stability and show a high temporal coherence with these reference signals. These EVM-based heart rate estimates also show a higher similarity with ECG in monkeys than in human subjects (max difference in BPM: human: 8.28 BPM; monkeys: 3.91 BPM) probably due to the fact that monkeys were under anaesthesia while human subjects were awake. Irrespective of anaesthesia and quite remarkably, coherence between EVM-based and ECG-based heart rate estimates tended to be higher when extracted from monkey videos than from humans. While this could still be due to improved stability of the face in the video stream in the anesthetized preparation, it still noteworthy, as human faces have more glabrous skin as well as a thinner skin, thus allowing for a better capture of superficial vascular changes. In any case, reporting this temporal coherence measure is crucial as it indicates that EVM-based heart rate estimation is sufficiently sensitive to track actual physiological variation of heart rate in time. These EVM-based heart rate estimates compare to those reported in a previous study extracting monkey heart rate from videos using imaging photoplethysmogram (Unakafov et al., 2018). Importantly, and in contrast with this previous study, EVM-based heart rate estimates can be reliably obtained both from RGB and IR videos.

Our present work addresses one crucial aspect of highest relevance to behavioural and cognitive studies in non-human primates, namely heart rate estimation in awake behaving monkeys. Here, using our EVM-based heart rate estimation method, we reproduce the observation that emotional sensory stimuli, whether it be auditory or visual, induce heart rate changes in monkeys (Bliss-Moreau et al., 2013). Reported heart rate changes can thus be specifically associated to stimulus emotional informational content rather than to visual stimulus presentation. This is an important point as local luminosity changes due to for example visual stimuli presentation have been shown to affect plethysmographic signals (Lewandowska et al., 2011) but not ambient light (Y. Sun et al., 2012). Indeed, heart rate has been shown to be modulated in a variety of tasks: emotional or valence tasks (Bauer, 1998; Bliss-Moreau et al., 2013; Kreibig, 2010; Lang, 1995), attentional tasks (Duschek et al., 2009; Gazzellini et al., 2016; Petrie Thomas et al., 2012; Siennicka et al., 2019) as well as learning tasks (Zeki Al Hazzouri Adina et al., 2014). Tracking this physiological parameter non-invasively, is methodologically easier, brings no stress and thus improves animal welfare while experimenting, saves training time and minimizes possible measure biasing factors (for example when animals are training to accept pulse oximetry measures). In addition, heart rate measure has been shown to covary with pupil dilation, typically controlled by the autonomic nervous system (Janig and McLachlan, 2013). Pupil dilation has been shown to vary as a function of reward (Satterthwaite et al., 2007; Varazzani et al., 2015), surprise (Lavin et al., 2014), vigilance for social distractors (Ebitz et al., 2014), arousal (Kennerley & Wallis, 2009) and systemic pharmacological neuromodulation (Reynaud et al., 2019). Last, heart rate can be a crucial parameter to assess behaviour following a drug induction (Bloch et al., 1973) or a lesion (Mitz et al., 2017). Overall, this ability to track changes in heart rate measure in an experimental context, that we demonstrate here, is thus of major implications to behavioural, cognitive and neuroscience experiments in awake macaque monkeys. This method is less expensive and requires no training. It can thus now be recommended and implemented as a control measure in most such awake macaque experiments.

In order to improve heart rate detection from videos, a recent paper suggests to use neural networks to detect and enhance the ROI from which the signal is extracted (Pursche et al., 2019). This technique can be implemented to detect moving animals in open fields or working animals in less restrained environments than ours, provided a high-resolution fast frame rate camera is used. This would be an important add-on to our work as heart rate and heart rate variation in monkeys in their home cage, during their time in the experimental setup or in open zoo spaces is considered as a good health state and stress marker (Clarke et al., 1994; Grandi & Ishida, 2015; Tatsumi et al., 1990). For example, Hassimoto et al. (2004) tracked the first 3 months of acclimation of rhesus monkeys in a new laboratory environment and they demonstrate a decrease of heart rate following this stressful period (Hassimoto et al., 2004).

Overall, we thus believe that the method we describe here is an easy and reliable non-invasive alternative for heart rate estimation both in anesthetized and awake monkeys. It is very easy to set up as it only requires a decent recording camera and a computer to implement the processing pipeline. In addition, it does not require any specific training from the animal. It can thus be applied during control veterinary or health visits under sedation or light anaesthesia, or under minimal contention condition in awake monkeys, or possibly even in home cage or open field environments.

The generalization of this method to experimental and ethology animal facilities can be considered as a contribution to animal welfare. During actual cognitive, behavioural or neuroscience experiments it can be considered as a refinement measure as well as a crucial physiological control parameter to include in the data analyses or in the framework of data sharing consortia (Margulies et al., 2020; Milham et al., 2018). By extension, we expect that this method can also be applied on young infants in neurodevelopment studies, where pulse oximetry heart rate tracking can turn out to be extremely challenging.

## Contributions

Conceptualization, S.B.H. M.F. and Q.G.; Stimuli preparation, M.H., M.F, Q.G, M.G; Data Acquisition, M.F. Q.G. M.G.; Methodology, Q.G., M.F. and S.B.H; Investigation, M.F., Q.G. and S.B.H.; Writing – Original Draft, M.F, S.B.H. and Q.G; Writing – Review & Editing, S.B.H., M.F., Q.G; Funding Acquisition, S.B.H.; Supervision, S.B.H.

